# Repression of Aurora kinase B prevents growth and tissue invasion in medulloblastoma

**DOI:** 10.1101/2023.06.23.545872

**Authors:** Alexandre Gries, Karthiga Santhana Kumar, Fabien Kuttler, Michael A. Grotzer, Martin Baumgartner

## Abstract

Novel treatment strategies are required to overcome therapy-associated sequelae in survivors of pediatric medulloblastoma (MB) while maintaining therapeutic efficacy. The tissue impact on drug response in MB is not well understood and drug profiling in the physiological context of the tissue may reveal novel therapy targets.

To gain insights into the growth and dissemination behavior of the MB tumor cells under treatment, we combined three-dimensional cell culture screening with *ex vivo* organotypic cerebellum slice co-culture (OCSC), which allowed assessing tumor cell behavior in the tissue context.

We screened a panel of 274 kinase inhibitors and identified aurora kinase B (AURKB) as a potential anti-invasion drug target in MB. We validated tumor suppressive activities of the AURKB inhibitor (AURKBi) barasertib and the structurally unrelated compound GSK-1070916 in cerebellum slice culture models for SHH, Grp3 and Grp4 MB at nanomolar concentrations. We confirmed the necessity of AURKB for tumor growth through genetic suppression of AURKB by siRNA in the tissue context. We revealed that the combination of AURKBi with the SRC/BCR-ABL inhibitor dasatinib acts synergistically to repress tumor growth and invasiveness in the SHH MB cell model ONS76, but not in Grp3 MB cells. Finally, we demonstrate that pharmacological repression of AURKB in the tissue context is as effective as X-ray irradiation to repress tumor growth.

Our data highlight that AURKBi is equally efficient as irradiation, suggesting that pharmacological targeting of AURKB may constitute a novel means to overcome radiotherapy limitations in patients younger than three years.

**Importance of the study:** Our data demonstrate anti-tumor activity of AURKB inhibitors in the tissue context for MB tumor cells that are *in vitro* resistant to the treatment. This indication of a tissue component to drug response contributes critical insights for drug response profiling for brain tumors. AURKB inhibitors barasertib and GSK-1070916 block growth and invasiveness of SHH, Grp3 and Grp4 MB tumor cells as well as primary ATRT. Importantly, AURKBi is equally efficient as irradiation. In conclusion, a tissue component contributes to sensitivity to AURKB inhibition and pharmacological targeting of AURKB may constitute a novel means to overcome radiotherapy limitations in patients younger than three years.

**Key points:**

- AURKB is essential for in tissue growth of MB.
- Inactivation of AURKB causes p53 upregulation and decreased in tissue growth and dissemination.
- AURKB inhibition in the tissue context is as effective as irradiation to restrict tumor cell growth.

## Introduction

Brain and nervous system tumors represent the second most common cancer entity in terms of incidence and mortality in children worldwide [Cancer Today, WHO], within which medulloblastoma (MB) is the most common malignant tumor. Current MB treatments consist of surgery complemented with chemotherapy and radiotherapy for children older than three years. Survivors often suffer from treatment-associated long term side effects, and approximately 30% of patients still die from the disease^1^. Four molecular subgroups (WNT, SHH, Group 3 (Grp3), and Group 4 (Grp4)) sub-divided into twelve subtypes have been molecularly characterized^2^. Although Grp3 and Grp4 MB tumors present a higher incidence of distal metastasis, metastasis is also a poor prognosis marker for SHH subgroup tumors^1–3^. MB metastasis occurs locally in the cerebellum midline and hemispheres and distally on the leptomeninges of the brain and the spinal cord. Local recurrence as a consequence of tissue invasion is observed for SHH MBs, whereas Grp3 and Grp4 tumors tend to spread more distally^1^. Growth factor signaling pathways have been implicated in cerebellar tissue invasion^4–6^ and chemokine signaling in distal spreading^7^. Druggable driver kinases of MB dissemination and distal growth remain to be identified.

Aurora Kinase A (AURKA), Aurora Kinase B (AURKB) and Aurora Kinase C (AURKC) are highly conserved proteins with homologous structure involved in mitosis^8^. AURKA controls centrosome maturation, bipolar spindle assembly, and cytokinesis, while AURKB and AURKC allow chromosome condensation, attachment to kinetochores and their alignment during metaphase and cytokinesis^9^. AURKA was identified as critical gene involved in MB progression^10^ and AURKA inhibition in MB cell lines was found to increase chemosensitivity^11^. Exposure of SHH MB models to pan-AURK inhibitors were reported to enhance the sensitivity to conventional chemotherapies *in vitro* and *in vivo*^12^. AURKB expression correlates positively with MYC in MB and specific inhibition of AURKB blocks MYC overexpressing MB tumors *in vivo*^13^.

The aim of this study was to identify druggable kinases involved in invasion and dissemination control in MB. We used the spheroid invasion assay (SIA)^14^ in combination with a panel of 274 pharmacological kinase inhibitors to identify kinases involved in invasion control in MB. Pan AURKA/B inhibitors identified AURKB as a potential druggable target in MB. Since no study so far systematically addressed specific pharmacological targeting of AURKB in MB in the tissue context, we validated AURKB inhibition *ex vivo* using the organotypic cerebellum slice co-culture (OCSC) model for MB tumors^15^. We further compared efficacy of pharmacological AURKBi with X-ray radiation, to explore AURKBi as a potential replacement treatment for children not eligible for radiotherapy.

## Materials and Methods

### Tumor cells

Culture, origin and authentication of ONS-76 (SHH), HD-MB03 (Grp3), D425-Med (D425, Grp3) and D283-Med (D283, Grp3/4) cell lines are described in supplementary file 1. Generation of LA-EGFP-lentivirus transduced descendants is described here:^5^.

### Patient-derived cells

Patient-derived tumor tissue samples were obtained from in-house patients with informed patient consent (Approval by: Cantonal Ethic Commission Zurich, Switzerland) and tumor tissue samples were processed as described^16^. 10’000 patient-derived cells (PDCs)/well were seeded in 96-well clear round bottom low adhesion plate to obtain spheroids, which were thereafter implanted on cerebellum slices. 300’000 PDCs were seeded in 6-well low adhesion plate for 48 h to extract RNA and perform RT-qPCR.

### Kinase inhibitor screen

1’200 individual samples containing 3’000 ONS-76 mCherry-nuc cells/well were seeded in low adhesion plate for 24h. Spheroids were embedded in collagen I as described in^14^, and treated individually with a total of 274 kinase inhibitors at 3 μM concentration for 24h. The sum of distances between each cell and spheroid were determined (see supplementary file 1 for details), and a z-score for each sample was used to establish a hit list of effective compounds^17^.

### Spheroid Invasion Assay (SIA)

SIA was performed as described in^14^ with 2’500 ONS-76 cells seeded in 100 μl per well of complete RPMI medium in 96-well low adhesion plate. Cell invasion was expressed as the sum of invasion distances of the cells from the center of the spheroid after 24h.

### Viability and Cytotoxicity assay

CellTiter Glo® and CellTox™ Green assays were used with 750 cells seeded in 25 μl per well of complete medium in low adhesion plates for 24h. Kinase inhibitors or DMSO for normalization were deposited on plates using a HP D300 Digital Dispenser (Hewlett-Packard Development Company). The relative luminescence (CellTiterGlow) or fluorescence (CellToxGreen, λ_ex_=485nm and measuring at λ_em_=520nm) per well was measured after 24, 48 and 72h of compound incubation at 37°C using a Cytation 3 imaging reader (BioTek®). ZIP (zero interaction potency) synergy score of drug combinations was obtained using SynergyFinder® (https://synergyfinder.fimm.fi/).

### Irradiation (IR)

X-ray irradiation of cells in suspension cultures or OCSCs was performed using the CellRad X-ray irradiator PXI (Precision, North Branford, CT, USA) with the following settings: 1,8 Gy, 0,6 Gy/min, 5 mA, 105 kV.

### Long term viability assay

15’000 cells in 60 μl cell culture medium were seeded on Millipore inserts. The inserts were then transferred to 24-well plate containing 500 μl of OCSC medium per well. Medium was changed daily, and cells were treated with 10, 100, or 500 nM barasertib for five consecutive days. Cell viability was determine by CellTiter Glo 3D assay.

### Immunoblot (IB)

200’000 cells were seeded in 6-well low adhesion plates for 48h in complete medium and all treatments were performed in suspension. Cells were lysed with laemmli buffer. For primary antibodies used see supplementary file 1. HRP-linked secondary antibodies were used with either Pierce™ ECL Substrate or SuperSignal™ West Femto Maximum Sensitivity Substrate. Integrated density of immuno-reactive bands was quantified using Image Lab (Version 5.2.1, Bio-Rad Laboratories).

### Organotypic cerebellum slice co-culture (OCSC)

OCSC cultures were authorized by the Cantonal Veterinary Service of Zürich (ZH116/20) and performed as described in^15^. A maximum of three slices were placed per insert, and media were changed daily for two weeks. Tumor spheroids from ONS-76 LA-EGFP, HD-MB03 LA-EGFP, D425, D283 or from primary tumor derived cells were then placed on the slices and incubated for 24 h. The tumor spheroid-slice co-cultures were subsequently drug-treated for five days.

Click-iT® EdU (C10340, Invitrogen) was used for detecting proliferating cells. 20 min EdU incubation was used for established tumor cell lines and 2h for patient derived cells. Antibodies used are listed in supplementary file 1. The inserts were flat mounted onto glass slides in glycergel mounting medium.

Images were acquired on a SP8 Leica confocal microscope (Leica Microsystems, Mannheim, Germany). Tumor volume (TV) and proliferation volume (PV) were determined by quantifying the volume of the green fluorescence (human nucleoli) and the volume of the red fluorescence (EdU staining), respectively, from z-stacks using Imaris software (volume thresholding: smooth surfaces detail 15 μm [human nucleoli] and 5 μm [EdU], background subtraction [local contrast] 8.52 μm). Two measurements were acquired per slices, which ere normalized to the negative control (DMSO-treated) condition (CTRL). Relative proliferation volume (rPV) = PV/TV.

### siRNA-mediated depletion

10 nM siAURKB was used for AURKB depletion using lipofectamine RNAiMAX transfection reagent with reverse transfection. 300 μl of the siRNA transfection mix were deposited in a 6-well plate onto which 300’000 MB tumor cells were seeded. After 6h of incubation, 5’000 transfected cells/well were seeded in a 96-well round bottom low adhesion plate for 24h to form spheroids, which were then deposited on cerebellum slices. Co-cultures were then maintained in culture for 48h (72h after transfection). Silencing of AURKB was validated 72h after transfection by IB from 250’000 cells seeded low adhesion plates.

### RT-qPCR: mRNA expression level after treatments

300’000 HD-MB03 or D425 were seeded per well in 6-well low adhesion plates. After 48 h, medium was replaced, cells were irradiated and incubated for another 8 or 24 h ± 50 nM barasertib. At endpoint, cells were collected and RNA was purified using QIAGEN RNeasy Mini Kit. 1 μg of mRNA was reverse transcribed in cDNA using the high-capacity cDNA Reverse Transcription Kit. RT-qPCR was performed using TaqMan Master mix under conditions optimized for the ABI7900HT instrument. The ΔCT method was used to calculate the relative gene expression of each gene of interest relative to *GAPDH*.

### Statistical analysis

Statistical differences were calculated with Kruskal-Wallis’ test followed by multiple comparison test with two-stage linear step-up procedure of Benjamini, Krieger and Yekutieli or with Mann-Whitney test with a p-value corrected using Prism GraphPad 8 software. We considered as significant p-value p< 0.05.

## Results

### 3D invasion assay identifies Aurora and SRC kinase inhibitors as invasion inhibitors

We tested 274 different kinase inhibitors in the SHH ONS-76 MB cancer cell line^18^ to identify inhibitors of pro-invasive kinases in MB. 60 inhibitors including dasatinib^19^ used as positive control reduced ONS-76 cell invasion (Fig. 1A, supplementary table 1). The top 20 compounds included several aurora kinase inhibitors (AURKi) (Fig. 1A,B). To validate the potential implication of SRC and AURK, we selected SRCi ponatinib, saracatinib, and dasatinib and AURKi Aurora kinase A inhibitor I (AURKAi I), tozasertib (VX-680), and TAK-901. We first confirmed the specific inhibition of SRC and AURKs by immunoblot (IB) using phosphosite-specific antibodies^20,21^ (Fig. 1C). The SRCi inhibited SRC kinase activity at 1 μM concentration without affecting AURK activities. We also confirmed AURKAi I specificity for AURKA. Tozasertib and TAK-901 reduced both AURKA and AURKB phosphorylation. All three SRCi effectively repressed collagen I invasion of ONS-76 cells at nanomolar concentrations (Fig. 1D, Supplementary Fig. 1A), consistent with the EC50 of dasatinib in this line^19^. TAK-901 suppressed invasion of ONS-76 cells at 50 nM concentration. Tozasertib reduced collagen cell invasion in a dose dependent manner with an EC50 of approximately 4 μM, whereas AURKAi I did not repress invasion up to 20 μM concentration (Fig. 1D). AURKAi I is a specific inhibitor of AURKA, tozasertib and TAK-901 could also target AURKB (Fig. 1C). The two AURKB inhibitors^22^ barasertib (AZD-1152) and GSK-1070916, which both effectively reduced AURKB phosphorylation without affecting AURKA (Fig. 1C), were also ineffective in reducing collagen I invasion (Fig. 1D, Supplementary Fig. 1A). We therefore combined AURKAi I with either barasertib or GSK-1070916 to test whether the inhibition of both AURKA and AURKB is necessary for blocking invasiveness (Supplementary Fig. 1B-C). As neither combination showed a reduction in ONS-76 cell invasion, we concluded that TAK-901 and tozasertib effects are likely pan-kinase off-target activities of these inhibitors. None of the tested inhibitors at effective concentration reduced cell viability (Fig. 1F) or induced cell death (Fig. 1E)

**Figure 1:**
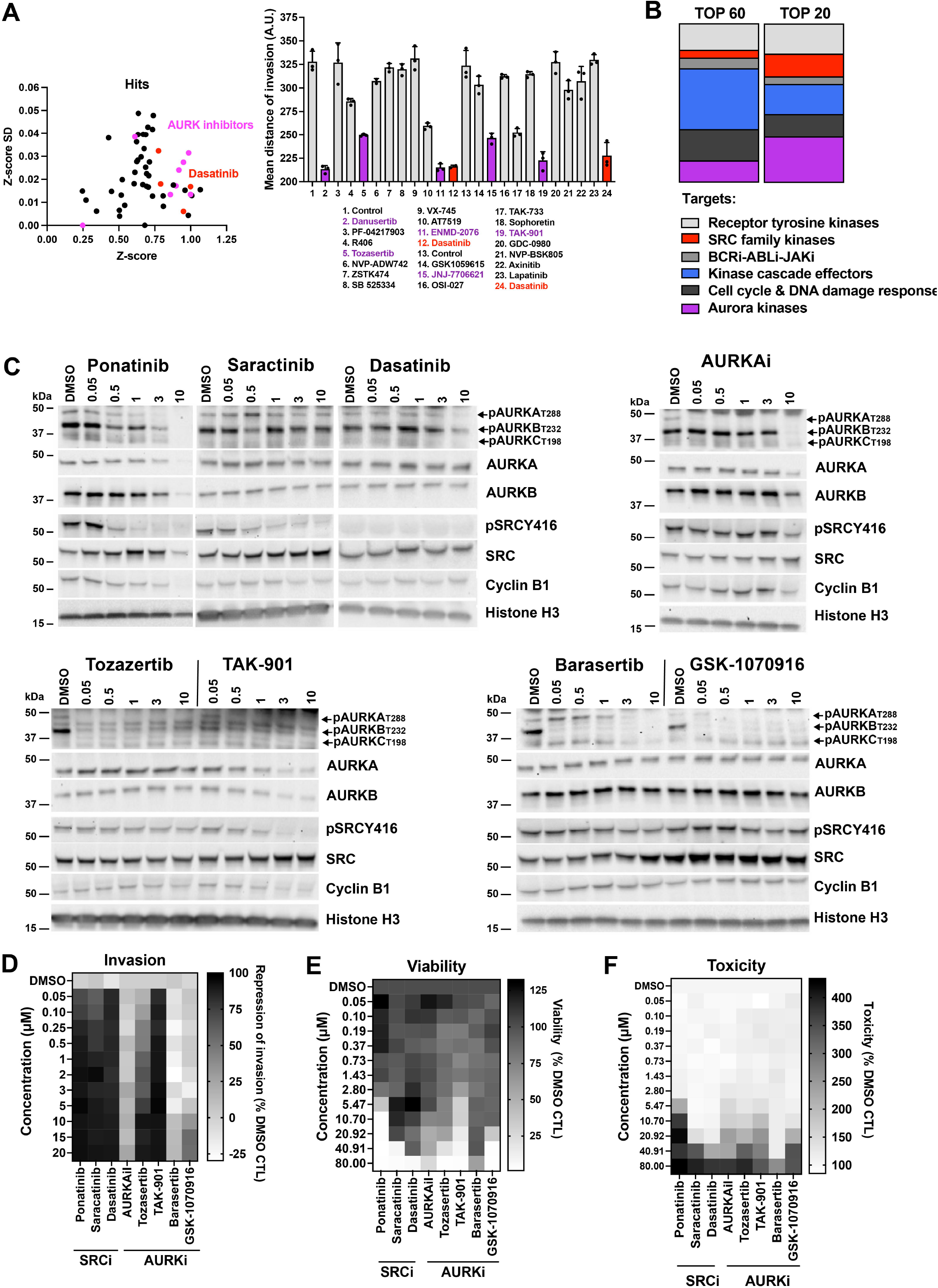
Blockade collagen I invasion by Aurora and BCR-ABL/SRC kinase inhibitors. **A)** Left: X/Y plot of Z-score and Z-score SD of hit list. Right: Validation of selected drugs using aCDc software^14^ showing mean distances of invasion from center of spheroids and SD (n=3 biological replicas). Purple: AURKBi, red: BCR-ABL/SRC inhibitors, all tested at 3 μM. **B)** Representation of top 60 and top 20 candidates across kinase inhibitor families. **C)** IB using antibodies specific for the phosphorylation of the active sites of SRC, and aurora kinase A and B proteins. Cyclin B1 protein expression confirms cell proliferation. **D)** Heat-map of mean values of anti-invasion efficacy of selected kinase inhibitors of SIA shown in A). **E)** Heat-map of mean cell viability in ONS-76 cells. **F)** Heat-map of cytotoxicity in ONS-76 cells. (D – E: n=3 biological replicas).

In conclusion, we found that ponatinib, saracatinib, dasatinib, tozasertib and TAK-901 possess excellent anti-invasion efficacy at concentrations below 1 μM. In contrast, specific inhibition of AURKA and AURKB by AURKAi I and barasertib or GSK-1070916, respectively, is not sufficient to block collagen I invasion in ONS-76 cells *in vitro*.

### AURKB inhibition reduces growth and dissemination in the tissue context

We next tested a 5 days treatment regimen at 500 nM concentration on tumor cell growth and dissemination in the tissue context using OCSCs^15^ (Supplementary Fig. 2A). We calculated the DMSO-normalized tumor volume (TV) and relative proliferation volume (rPV) for each compound. Both saracatinib and dasatinib, reduced TV and rPV of ONS-76 cells at day 6 by 45% (p=0.0338) and 60% (p=0.0003), respectively (Fig. 2A, Supplementary Fig. 3A, 4 A,B). We observed no inhibitory effect of ponatinib. Barasertib and GSK-1070916 reduced ONS-76 TV and rPV significantly in OCSCs by around 40-50% (p≤0.0338) (Fig. 2A, Supplementary Fig. 3A, 4A,B,E). The TV of the Grp3 line HD-MB03 increased 5-fold between day 1 and day 6 under control (DMSO) conditions (Supplementary Fig. 3B, 4C,E). The SRCi showed little effect on TV and rPV in HD-MB03 cells (Fig. 2A, Supplementary Fig. 3B, 4 C-E). In contrast, both barasertib and GSK-1070916 significantly reduced TV and rPV by >50% (p≤0.0051) and 80%-90% (p≤0.0002), respectively (Fig. 2A, Supplementary Fig. 3B, 4C-E). In ONS-76 cells, the combination of SRCi and AURKBi led to the near complete eradication in the tissue context (Supplementary Fig. 3A, 4A,B,E). This additive effect of compound combination was less pronounced in the HD-MB03 cells, as AURKBi alone was already highly effective (Supplementary Fig. 3B, 4C-E).

**Figure 2:**
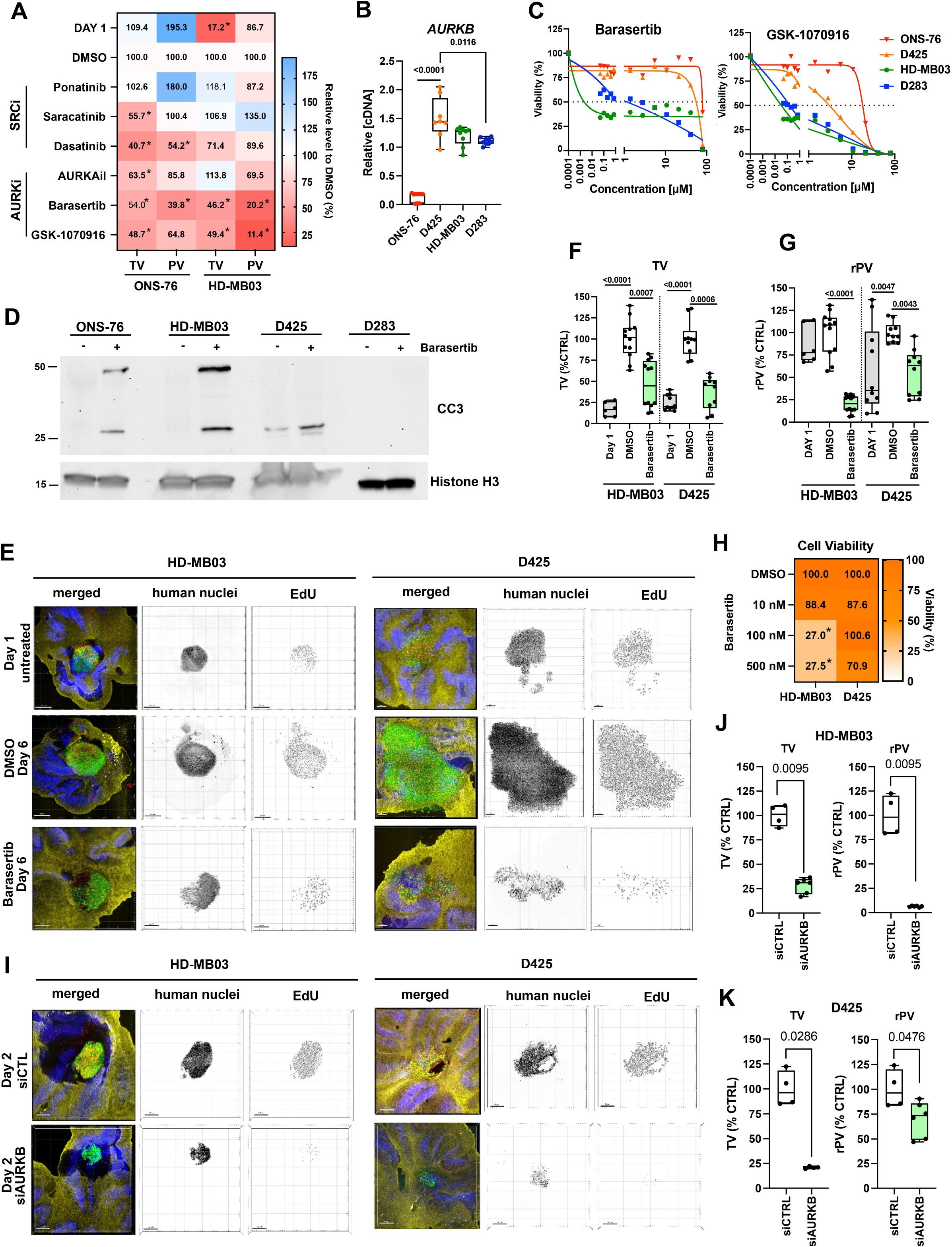
AURKBi reduces medulloblastoma cell dissemination *ex vivo*. **A)** Heat-map of tumor volume (TV) and relative proliferation volume (rPV = PV/TV) of ONS-76 and HD-MB03 cells in OCSCs after 5 days of treatment (*<p0.0338). **B)** RT-qPCR analysis of *AURKB* expression (n=3 biological replicas). **C)** Relative cell viability in response to barasertib and GSK-1070916 (n=3 biological replicas). **D)** IB analyses of anti-cleaved caspase 3 (CC3) and anti-histone H3 of cells treated with 0.05 (HD-MB03) or 3 (ONS-76, D425, D283) μM barasertib. **E)** Confocal microscopy images of OCSCs with implanted HD-MB03 and D425 cells (green). OCSCs with implanted tumor cells were treated with barasertib at either 500 nM (HD-MB03) or 100 nM (D425) concentrations. Red: Click-iT® EdU, blue: anti-calbindin, and yellow: anti-GFAP. **F)** TV of n=5 individually implanted spheroids. **G)** rPV of n=5 individually implanted spheroids. **H)** Heat map of *in vitro* cell viability of HD-MB03 and D425 cells after 5-days of treatments (n=3 biological replicas, *p<0.0018). **I)** Confocal microscopy images of OCSCs with implanted HD-MB03 and D425 cells (green) with siRNA-mediated depletion of AURKB in tumor cells. **J)** TV of experiment shown in I. Data of n=3 individually implanted spheroids are shown. **K)** rPVs of experiment shown in I. Data of n=3 individually implanted spheroids are shown.

The Grp3 cell lines D425 and HD-MB03 as well as the Grp3/4 D283 cell line display considerably higher *AURKB* expression than SHH ONS-76 cells (Fig. 2B). Similar relative expression of AURKB is also observed at protein level, although D283 cells express less AURKB than HD-MB03 and D425 (Supplementary Fig. 5A). Barasertib effectively reduced viability of HD-MB03 and D283 cells *in vitro*, with an IC50 of 2 nM and 1 μM, respectively, unlike the D425 and ONS-76 cells, where we observed no inhibition up to 40 μM (Fig. 2C, Supplementary Fig. 5B). GSK-1070916 also effectively reduced HD-MB03 and D283 cell viabilities with IC50 of 0.033 and 0.2 μM, respectively. Barasertib treatment increased the cleaved caspase 3 (CC3) signals in HD-MB03, D425 and ONS-76 cells (Fig. 2D). No increase in CC3 was detected in D283 cells, suggesting that decreased viability of this line (Fig. 2C, Supplementary Fig. 5B) is rather the consequence of reduced proliferation and not of apoptosis.

To explore the therapeutic potential of AURKBi for Grp3 MB, we compared the effect of AURKBi in HD-MB03 and D425 cell lines in OCSCs (Fig. 2E-G). We selected these two cell lines because (i) they highly express MYC (Supplementary Fig. 5A), a known marker of MB aggressiveness^3^, and (ii) they represent an *in vitro* barasertib-sensitive (HD-MB03) and -resistant cell line (D425). D425 cells display a similar propensity to grow and invade the cerebellum tissue as HD-MB03 (Fig. 2E). Barasertib treatment significantly reduced TV in HD-MB03 and D425 by 64% (p=0.0007) and 73% (p=0.0006), respectively. Similarly, rPV was reduced in both cell lines with a reduction by more than 80% (p<0.0001) for HD-MB03 and by 45% (p=0.0043) for D425. To assess slice impact on treatment efficacy, we cultured HD-MB03 and D425 cells in the absence of the cerebellum slices on inserts for five days using the OCSC medium (Fig. 2H). Under these conditions, barasertib treatment reduced HD-MB03 viability by 70% (p=0.0018), similar to what we observed in the tissue context (Fig. 2B). In contrast, no significant reduction of cell viability was observed for D425 cells, whereas the same concentration of barasertib efficiently reduced TV and rPV of D425 cells in OCSCs (Fig. 2E-H). These results indicate that the cerebellar microenvironment can impact the functional activity of barasertib. AURKB-depletion in tumor cells by siRNA phenocopied barasertib effects and led to a near complete eradication of the tumor cells in OCSCs (Fig. 2I, 2J,K, Supplementary Fig. 6), thus confirming the relevance of AURKB function in the tumor cells for growth and dissemination of MB in the tissue context.

### Barasertib treatment increased p53 levels and induces p53 target gene expression

AURKB phosphorylation of MYC on S67 can stabilize MYC protein and control its transcriptional activity^23^. We did not observe a clear effect of barasertib treatment on MYC protein levels (Fig 3A), neither did we observe a reduction of *MYC* or of the MYC target genes *GLUT-1*^24^, *ITGA1*^25^ and *CDC42*^26^at mRNA level (Supplementary Fig. 7). AURKB can also phosphorylate p53 at S183, T211, and S215 to accelerate its degradation and suppresses target genes such as *p21*^27^. 50 nM barasertib is sufficient to repress AURKB Thr232 phosphorylation within 6h in HD-MB03 and D425 cells (Fig. 3A) and to increase p53 protein and CC3 levels within 24 h (Fig. 3B). Consistently, we also observed the induction of the p53 target genes *p21* and *BAX* (Fig. 3C-F). We used X-ray irradiation (IR) as control for p53 induction^28^, which caused p53 target gene expression within 8h. We did not observe a comparable induction of *p21* and *BAX* after 8h of barasertib treatment, suggesting that p53-induction is rather a long-term effect of AURKB inhibition.

**Figure 3:**
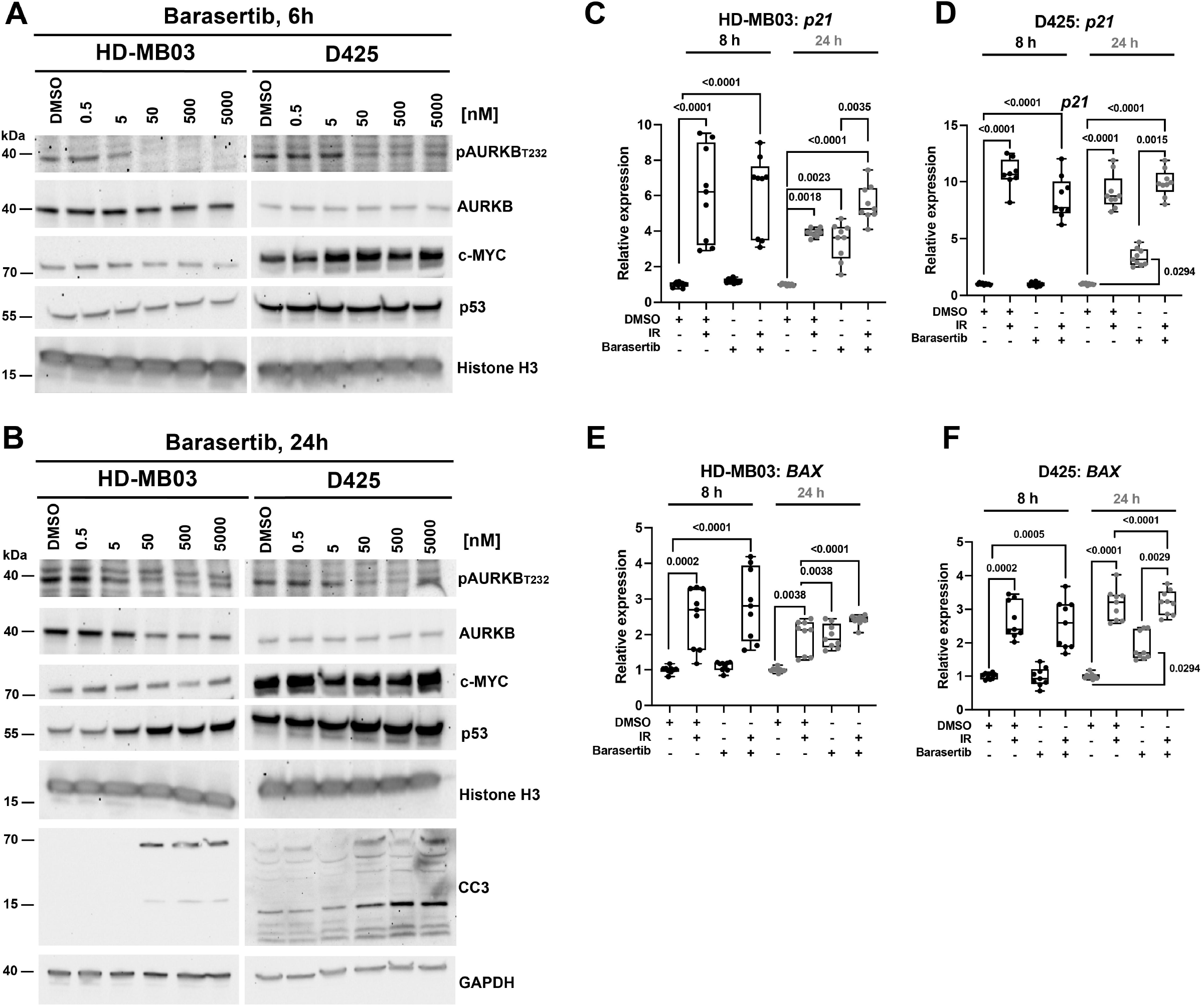
AURKBi induces p53 expression and CC3 in Grp 3 MB. **A)** IB of barasertib treatment effects on AURKA,B,C phosphorylation (pThr232) and MYC and p53 protein levels after 6h of treatment. **B)** IB of barasertib treatment effects on AURKA,B,C phosphorylation and MYC, p53 protein levels and CC3 after 24h of treatment. **C-E)** RT-qPCR quantifications of *p21* and *BAX* mRNA levels in HD-MB03 (B-C) and D425 (D-E) cells. Cells were irradiated (IR, 1,8 Gy) and/or treated with barasertib (50 nM) for 8 and 24 h (n=3 biological replicas).

Thus, reduced tumor expansion and proliferation after AURKB inhibition is not a consequence of impaired MYC function in MB but rather involves the induction of p53 target genes.

### Comparable efficacy of barasertib and IR in the tissue context

To explore whether barasertib could complement current IR replacement protocols^29,30^, we compared barasertib and IR treatment effects on cell viability. We first treated HD-MB03 and D425 cells *in vitro* with barasertib and/or IR (1,8 Gy) as depicted in Fig. 4A. In HD-MB03, as little as 20 nM barasertib reduced viability by more than 75% (Fig. 4B, Supplementary Fig. 8A). The same concentration had no effect in D425 (Fig. 4B, Supplementary Fig. 8B). IR reduced cell viability in both HD-MB03 (minus 55-63%) and D425 (minus 42-55%). Adding a second cycle of IR did not significantly increase treatment efficacy (Fig. 4A,B). The combinations of IR plus barasertib did not further reduce cell viability compared to IR or barasertib alone (Fig. 4A,B). In HD-MB03, barasertib treatment alone displayed a 25% (p=0.0018) increased inhibitory activity compared to IR treatment alone.

**Figure 4:**
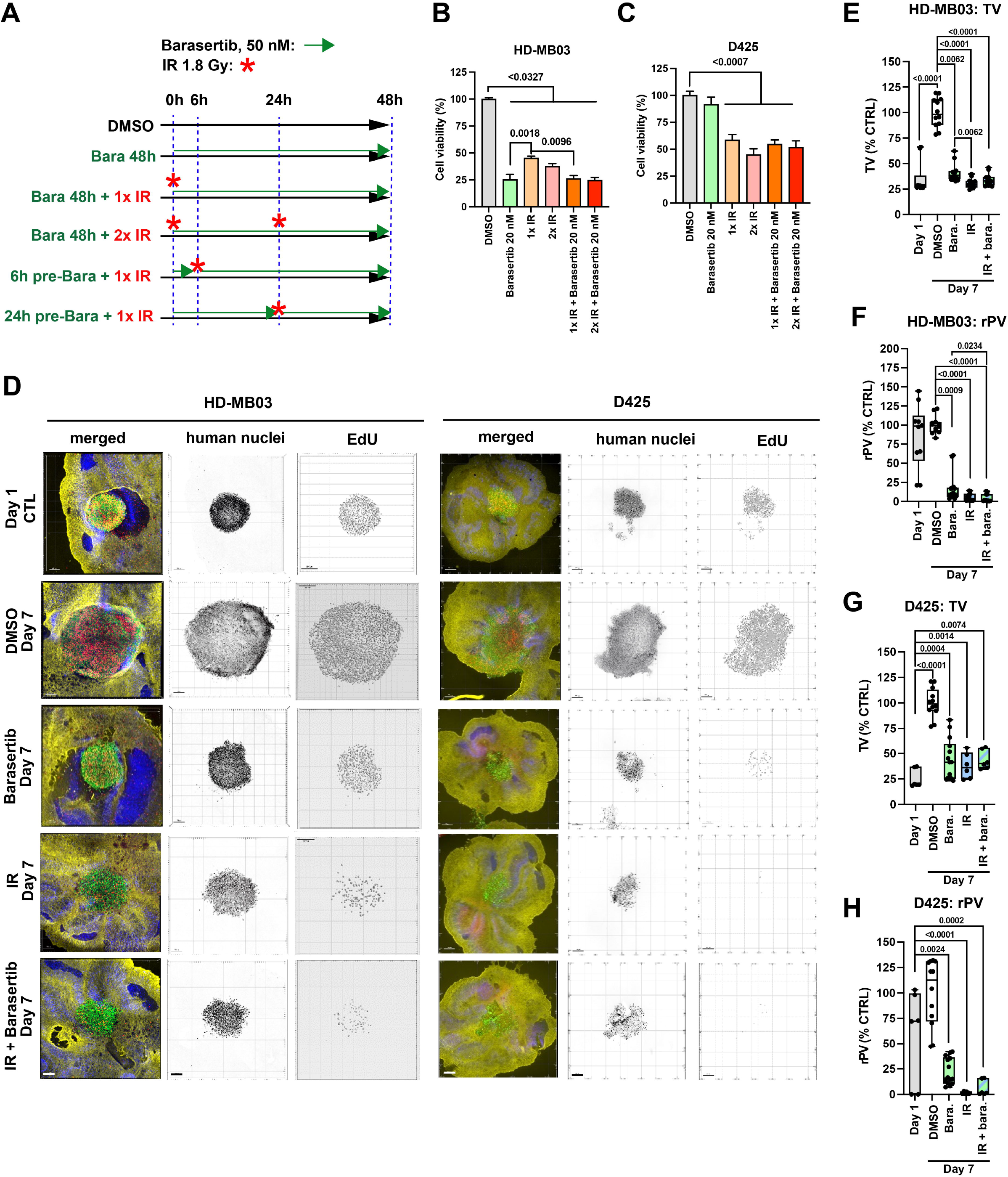
Comparable reduction of tumor growth in the tissue context after AURKBi or IR. **A)** Experimental procedure for the *in vitro* cell viability assays shown in B and C. **B)** Viability of HD-MB03 and **C)** D425 cells in response to IR (1,8 Gy) and barasertib treatment alone or in combination after 48 h (n=3 biological replicas). **D)** Confocal microscopy analysis of OCSCs 6 days after treatments with barasertib 100 nM and IR 1.8 Gy. Green: Anti-human nucleoli antibody, red: Click-iT® EdU, blue: anti-Cabindin Click-iT® EdU, yellow: anti-GFAP. **E-H)** Box plots of TV and rPV of OCSCs from D. (n≥3 biological replicas).

*Ex vivo* in OCSCs, IR reduced TV of HD-MB03 cells by 70% (p≤0.0001), and rPV by 95 (p≤0.0001) (Fig. 4D-F). In the same experiment, barasertib reduced TV by 60% (p=0.0062) and rPV by 83% (p=0.0009). The combination IR plus barasertib reduced TV by 66% (p≤0.0001) and rPV by 96% (p≤0.0001) (Fig. 4D-G). In D425 cells, IR reduced the TV by 63% (p=0.0014) and the rPV by 99% (p=0.0033), Barasertib reduced TV by 66% (p=0.0004) and rPV by 88% (p=0.0024) (Fig. 4D,G,H). The combination of barasertib and IR reduced TV and rPV in D425 by 57% (p=0.0074) and 94% (p=0.0002), respectively. In both cell lines, the combination of barasertib and IR did not significantly increase efficacy compared to each treatment alone. Together, our results indicate that AURKBi could represent a potential IR replacement treatment.

### Subgroup-specific sensitivities to combinatorial treatments in the tissue context

We next explored whether the combination of dasatinib and barasertib or GSK-1070916 could exert an additive or synergistic effect in HD-MB03 (Fig. 5A-B), D425 (Fig. 5C-D), ONS-76 (Fig. 5E-F) and D283 (Fig. 5G-H) cells. The responses differed considerably between the cell lines. In HD-MB03 cells, the barasertib-dasatinib combination elicits antagonistic (Fig. 5A-B) and in ONS-76 (Fig. 5E-F) or D283 cells (Fig. 5G-H) synergistic effects. We also noted some difference when comparing barasertib-dasatinib and GSK-1070916-dasatinib combinations, indicating that the difference in chemical structure of the two AURKBi results in a differential drug response.

**Figure 5:**
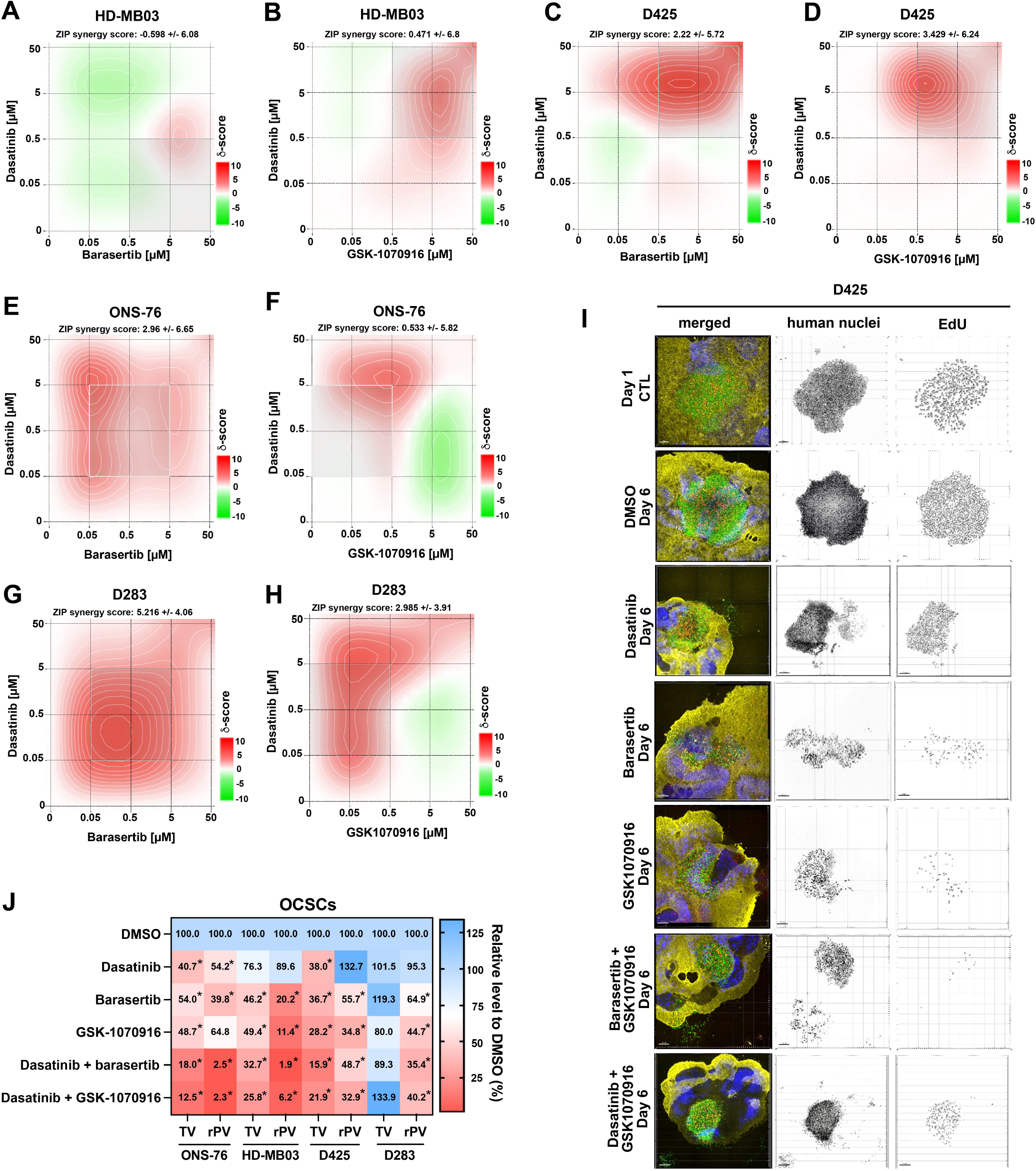
Synergistic activity of AURKBi and dasatinib *in vitro* and *ex vivo* A-H) Synergy score of drug combinations after 48 h of treatment with 0.005, 0.05, 0.5, 5 and 50 μM of barasertib and dasatinib in HD-MB03 **(A)**, D425 **(C)**, ONS-76 **(E)** and in D283 **(G)** cells. Synergy score of drug combination of GSK-1070916 and dasatinib in HD-MB03 **(B)**, D425 **(D)**, ONS-76 **(F)** and in D283 **(H)** cells. **I)** Confocal IFA of OCSC-embedded D425 cells treated for 5 days with 100 nM barasertib, GSK-1070916, or dasatinib, and combinations of barasertib and dasatinib or GSK-1070916 and dasatinib. Green: Anti-human nucleoli antibody, red: Click-iT® EdU, blue: anti-Cabindin, yellow: anti-GFAP. **J)** Heat map of TV and rPV of ONS-76, HD-MB03, D425 and D283 OCSC co-cultures after 5-days of treatments (n≥3 biological replicas, *p<0.0338).

We also evaluated the combinatorial effect of AURKBi and dasatinib, both at 100 nM concentration, in OCSCs with D425 (Fig. 5I, Supplementary Fig. 9D,E) and D283 (Supplementary Fig. 9A-C) cells. AURKBi strongly reduced TV and rPV in D425 (Fig. 5J), but not in D283 cells, where only rPV was reduced. Combining AURKBi with SRCi decreased TV of the ONS-76 cells by a factor of >-3 (p≤0.0478) and rPV by a factor >-17 (p=0.0300) compared to AURKBi single treatment alone. No significant additive effect of the compound combinations compared to AURKBi single treatment alone was noted in HD-MB03 cells. In D425 cells, we observed an additive effect of the barasertib-dasatinib combination on TV only (factor -2.3, p=0.0112) and in D283, we observed an additive effect of the barasertib-dasatinib combination on rPV only (factor -1.8, p=0.0139). Thus, AURKB plus BCR-ABL/SRC inhibition is additive for the SHH subgroup cell line ONS-76 only.

### Barasertib-dasatinib combination treatment blocks growth of primary tumor samples in the tissue context

We then evaluated the efficacy of barasertib alone or in combination with dasatinib in patient derived cells (PDC) from primary medulloblastoma (MB), atypical teratoid rhabdoid tumor (ATRT) and ependymoma (EPD) in OCSCs (Fig. 6A-B). In MB PDCs, the combination of barasertib and dasatinib reduced TV by nearly 59% and rPV by 56%. With this treatment, a similar level of repression of TV and rPV was also observed in PDCs of ATRT (-59%) and EPD (-49%) (Fig. 6A,B). In ATRT, we additionally tested single barasertib and dasatinib treatments. Barasertib decreased TV by 70% and rPV by 86%. No additive or synergistic effect was observed when barasertib was combined with dasatinib.

**Figure 6:**
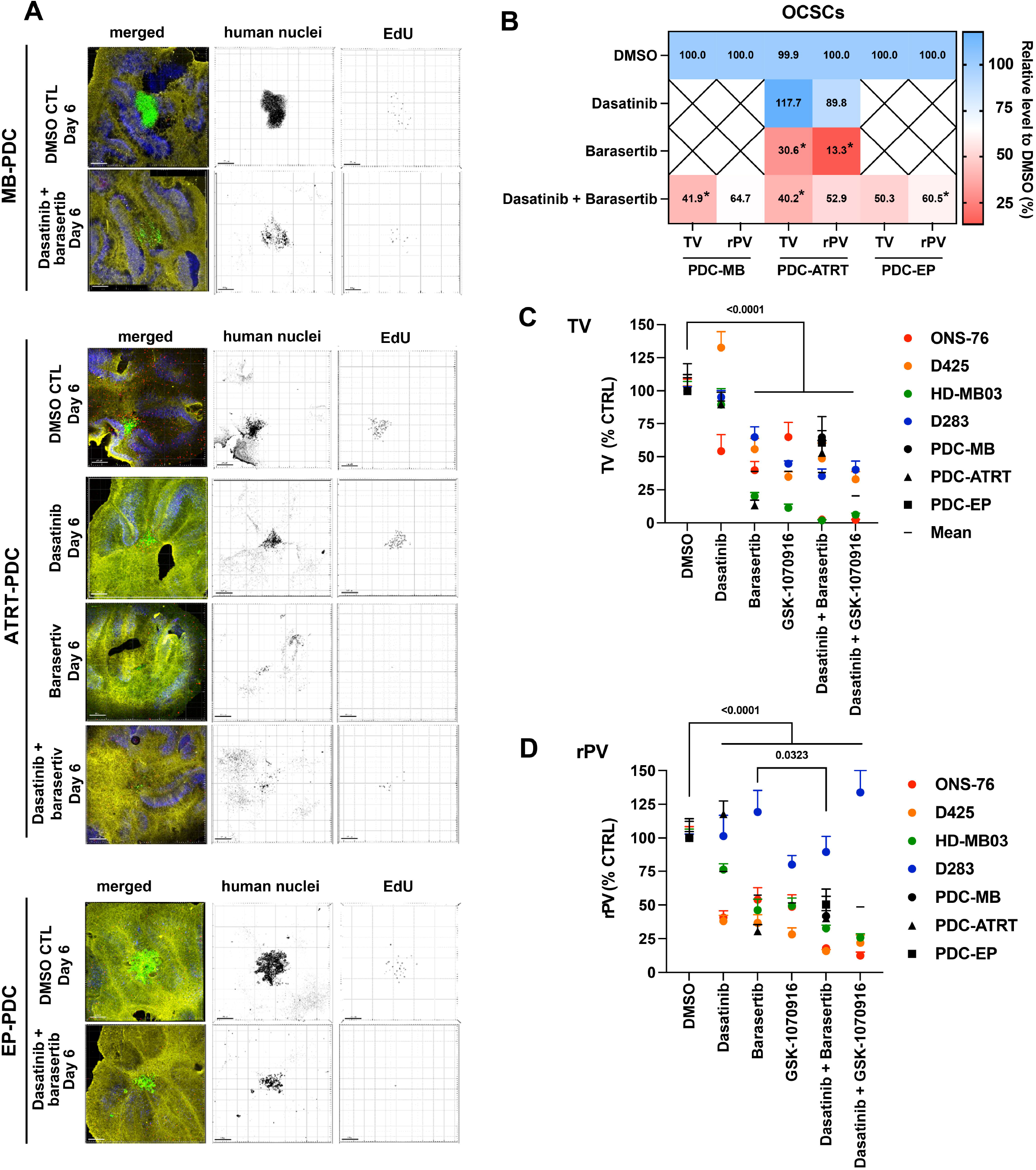
Barasertib effectively blocks patient derived cell growth in the tissue context. **A)** Confocal IFA of OCSC implanted MB-PDC, ATART-PDC and EP-PDC 5 days after treatments with 500 nM barasertib or dasatinib or the combination of barasertib and dasatinib. Green: Anti-human nucleoli antibody (PDCs), red: Click-iT® EdU, blue: anti-Cabindin, yellow: anti-GFAP. MB: medulloblastoma, ATRT: atypical teratoid rhabdoid tumor, EP: ependymoma. **B)** Heat map of TV and rPV of PDC-OCSC co-cultures after 5-days of treatments (n≥3 biological replicas, *p<0.05). C-D) TV and the rPV of all cell lines and PDCs used (Mean + SEM). **C**,**D)** Compilation of all TVs **C)** an rPVs **D)** from the OCSC experiments.

We also analyzed the drug response data of all OCSC experiments combined (Fig. 6C-D). AURKB inhibition by either barasertib or GSK-1070916 caused a significant reduction in TV and rPV in all cell lines and PDCs with the exception of D283 cells, where TV was not reduced. These data further corroborate the therapeutic efficacy of AURKB inhibition in restriction growth and tissue invasion in MB and – potentially – also in other high-grade pediatric brain tumors.

## Discussion

In the initial screen we used a 3 μM compound concentration. However, the invasion inhibition of some of the AURK inhibitors, such as tozasertib and TAK-901, observed at higher concentrations, is likely the result of off-target activity. TAK-901 at 3 μM concentration also repressed SRC Y416 phosphorylation, indicating that SRC kinase inhibition by TAK-901 could underly repressed invasiveness of ONS-76 at higher TAK-901 concentrations.

Drug exposure, drug import and drug extrusion mechanisms of tumor cells growing under 2D or 3D culture conditions differ from cells cultured as organoids or as xenografts *in vivo*^31,32^ and the cellular, chemical and biophysical context of the cerebellar microenvironment present *in vivo* is absent in current *in vitro* models. Some recent approaches addressed this problem and considerably improved 3D culture systems to better mimic MB tumor biology for high-throughput screenings^33,34^. Herein, we could demonstrate that despite the lack of *in vitro* efficacy, both barasertib and GSK-1070916 effectively reduced tumor volume and proliferation in the tissue context. This indicates a specific impact of the cerebellar microenvironment on the functional activities of drugs, analogous to the difference of drug sensitivity described recently in osteosarcoma^32^. Depletion of AURKB phenocopied the tumor inhibiting effect of barasertib and GSK-1070916, arguing against an indirect effect of barasertib on the cerebellar slice leading to tumor suppression.

Tumor cells expressing high MYC are sensitive to mitosis inhibitors^46^ and barasertib efficacy was demonstrated previously in a MYC-overexpressing D425 MB flank model *in vivo*^13^. In this study, 12 days after start of a four-day treatment, a considerable tumor re-growth was observed. This suggests that AURKB inhibition repressed tumor growth without eradicating the tumor cells. A significant reduction in tumor growth was also observed in a D425 intracranial tumor model^13^, where AZD-1152 (barasertib) peak levels of 0.7 ng/mg brain tissue were measured. This concentration corresponds to approximately ten-fold the effective concentration of 100 nM we observed in the cerebellar slices. To reach 0.7 ng/mg brain tissue, a roughly 140-fold higher concentration (mg/kg body weight) of barasertib was administered subcutaneously^13^. Although this demonstrated that effective concentrations of barasertib can be achieved in the brain, alternative routes of drug delivery may be necessary, such as intrathecal therapy via an Ommaya reservoir, which is widely used for the administration of chemotherapeutic drugs in brain tumors^37^.

To explore the suitability of the OCSC model for patient sample profiling and drug testing for primary pediatric brain tumors, we performed a pilot study using resection material from one primary MB, one primary ATRT and one primary EPD. For all three tumors, high suppressive efficacy of barasertib was demonstrated. Due to the limited material available, we are unfortunately not able to exclude the contribution of dasatinib to the tumor-repressive response of dastatinib-barasertib co-treatments in primary MB and EPD. However, in all experiments across cell lines we always observed superior efficacy of barasertib in comparison to dasatinib, and barasertib single treatment was highly effective in the ATRT. In an analogous approach, druggable vulnerabilities of brain metastasizing solid tumors were recently identified using in situ organotypic brain slice cultures^38^, further corroborating the clinical relevance of exploring drug efficacy *ex vivo* in the tissue context.

We confirmed barasertib efficacy in MYC overexpressing Grp3 MB^13^ in our cerebellar slice culture model and additionally demonstrate AURKBi efficacy in MYC-low expressing cell lines such as ONS-76. This indicates that AURKBi is effective in several MB subgroups and not only in MYC-overexpressing Grp3 tumors. Relevant transcription factors in aggressive MB are MYC/MYCN and p53^39–41^. We found no evidence that AURKB mediates its oncogenic functions through MYC. Our data rather supports a model where repression of AURKB leads to p53 stabilization and increased p53 target gene expression. Consistently, we observed upregulation of the p53 target genes *p21*^27^ and *BAX* after 24h of barasertib treatment, which also correlated with increased cleaved caspase 3.We expected increased p53 levels after barasertib treatment to sensitize the cells to IR^42^. However, both in HD-MB03 and D425 cells, the combination of AURKBi and IR did not significantly increase anti-proliferative or pro-apoptotic efficacy compared to each treatment alone. One possible explanation is that treatment-induced cell cycle arrest renders the cells less sensitive to either AURKBi or IR, which both target rapidly cycling cells. MB cell lines, primary tumors and xenografts express different isoforms of p53^43^, and depending of p53 basal expression levels, opposite roles of p53 on chemosensitivity of MB cells were described^44^. Determining the impact of AURKBi on differential p53 expression and function in MB will be necessary for better predicting patient’s responses. Importantly, our data demonstrate anti-tumor efficacy of barasertib similar to IR, which indicates that AURKBi could represent a potential replacement treatment strategy for IR therapy^29^. Finally, the increased treatment efficacy *ex vivo* in SHH MB cells when AURKBi and SRCi were combined implies a beneficial effect of co-repression of the previously described growth and invasion promoting activities of tyrosine kinase receptor signaling in this entity ^4,6^.

## Conclusion

We identified AURKB as a druggable vulnerability in medulloblastoma and demonstrate that two independent inhibitors of AURKB effectively repress MB tumor growth and dissemination in the cerebellar tissue. Our data further indicate that repression of AURKB is as effective as IR, suggesting that AURKB-targeting strategies could potentially replace or complement radiotherapy in young patients not eligible for radiotherapy.

## Authorship

Conceptualization: AG and MB; Methodology: AG, KSK, FK and MB; investigation: AG, KSK, FK and MB; access to human samples: MAG. Writing—review & editing: AG, FK, MAG, and MB.

## Data Availability

The data will be made available upon reasonable request

## Acknowledgments

The authors thank the Biomolecular Screening Core Facility of the EPFL and Jonathan Vesin and Julien Bortoli Chapalay for their competent technical support. We thank Ana Sofia Guerreiro Stücklin for providing patient samples and Sandra Laternser for technical assistance with patient samples. Imaging was performed with equipment maintained by the Center for Microscopy and Image Analysis, University of Zurich. Animals were kept in the infrastructure of the Laboratory Animal Services Center (LASC), University of Zurich.

