## Supplementary figures for "Repression of Aurora kinase B prevents growth and tissue invasion in medulloblastoma"

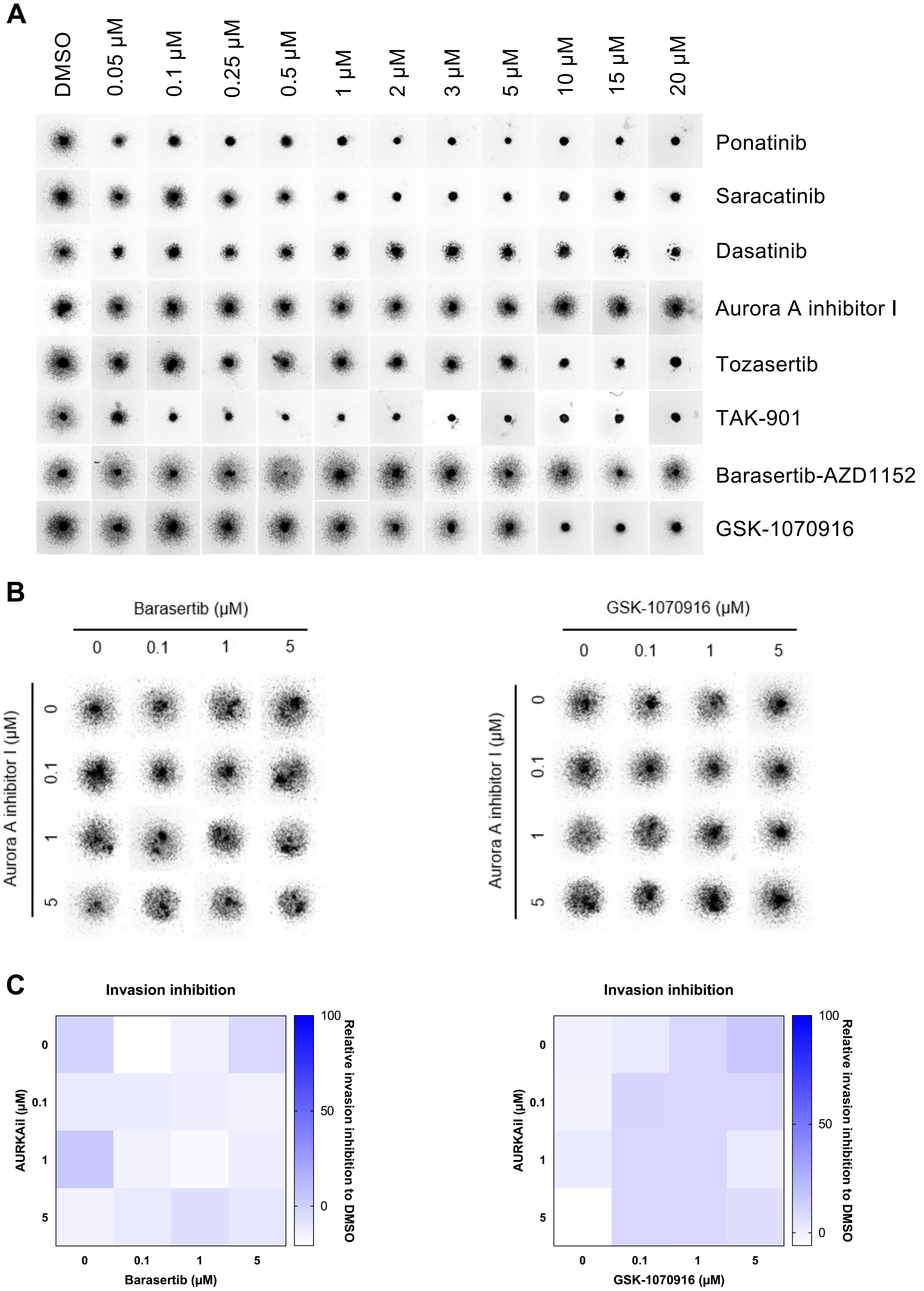


**Supplementary Figure 1: 3D spheroid invasion assay**

**A)** Representative pictures of the spheroid invasion assay (SIA), where cells, grown as spheroids, are embedded in collagen matrix +/- drug-treatment. Small spots surrounding the spheroids are the Hoechst-stained nuclei of the invading ONS-76 cells. Anti-invasion efficacy is presented in Figure 1D. **B)** Representative images of the SIA of ONS-76 cells treated with the combination of AURKA inhibitor with either of the two AURKB specific inhibitors Barasertib and GSK-1070916. **C)** Heat-map of triplicate measurements of SIA with ONS-76 cells treated as described in B. Darker blue indicates high anti-invasion efficacy.

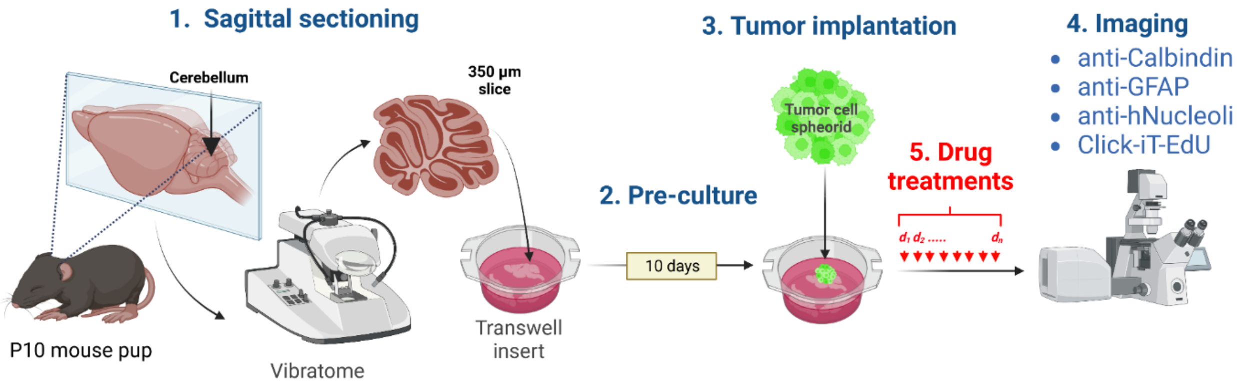


**Supplementary Figure 2: *Ex vivo* modelling of medulloblastoma tissue growth using organotypic cerebellum slice co-cultures (OCSCs).**

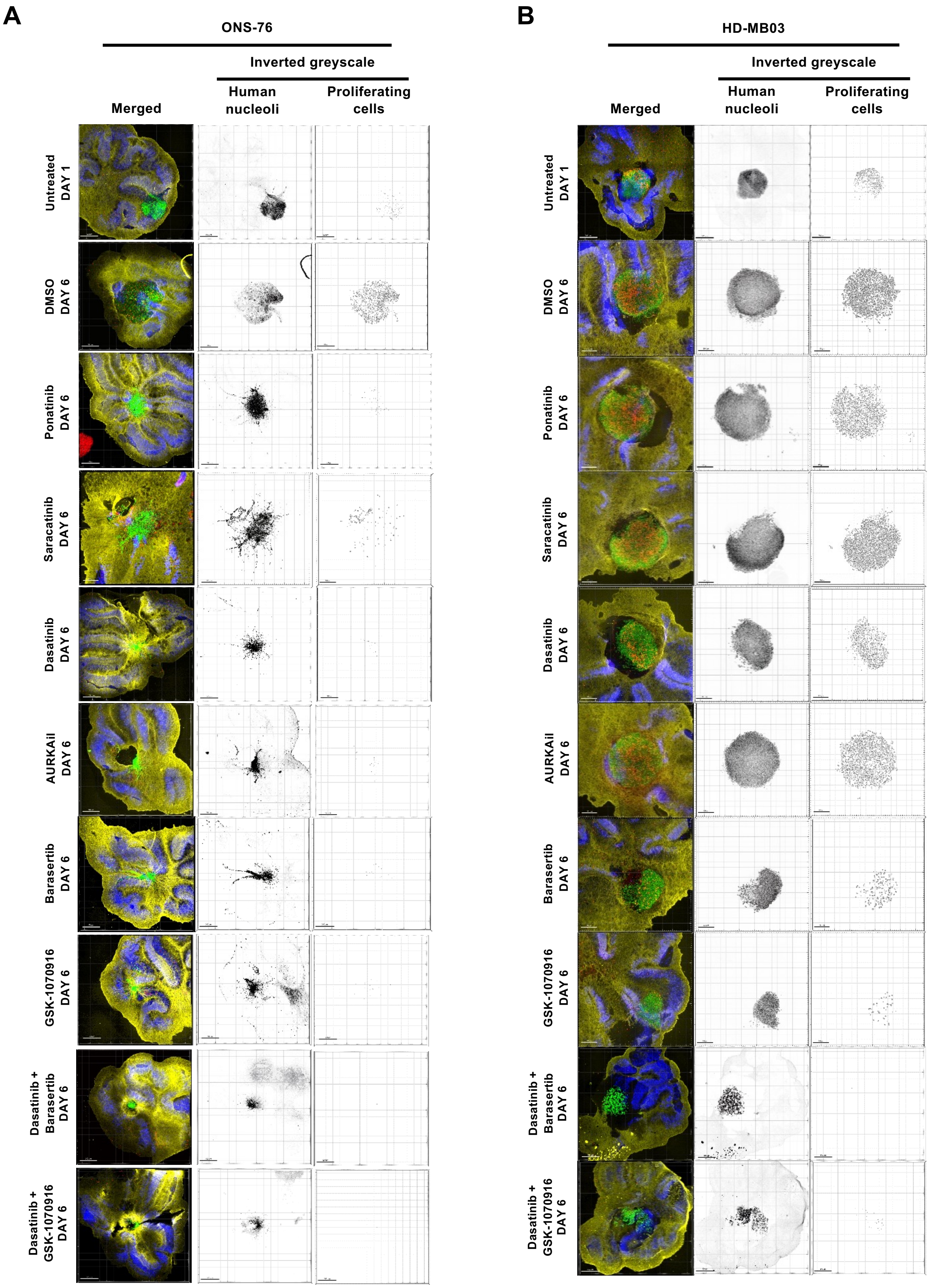


**Supplementary Figure 3: Representative images of OCSC experiment using ONS-76 and HD-MB03 cells**

Confocal microscopy images of OCSCs with ONS-76 **(A)** and HD-MB03 **(B)** cells (green) treated with indicated kinase inhibitors at 500 nM concentration. Red: Click-iT-EdU staining (proliferating cells), blue: Purkinje cells (anti-Calbindin), yellow: GFAP-positive cells (anti-GFAP). Quantifications are shown in Figure 2A, and in Supplementary Figure 4.

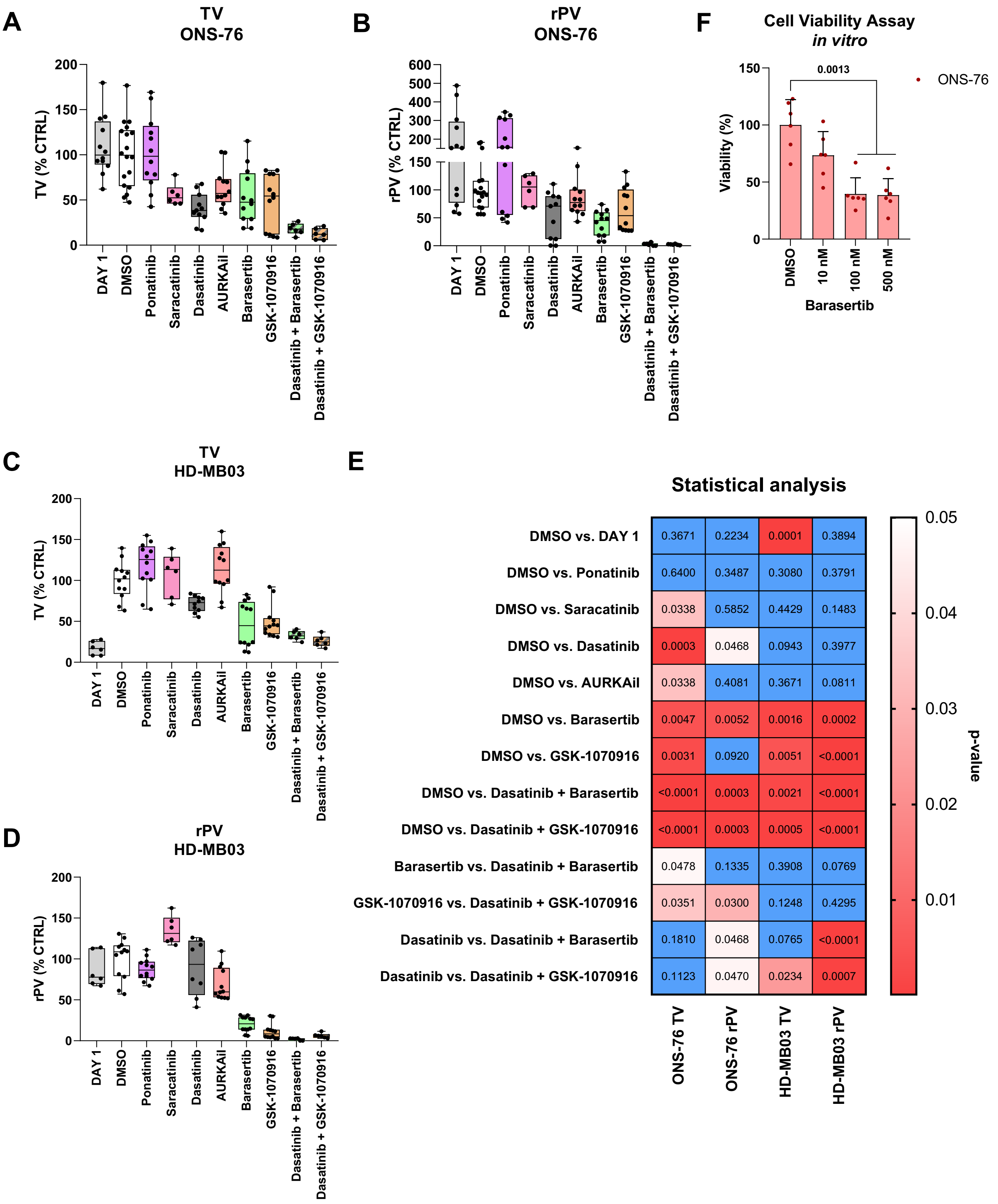


**Supplementary Figure 4: Quantification of OCSC experiments**

**A-D)** ONS-76 and HD-MB03 tumor cell volumes (TV) and relative proliferation volumes (rPV) relative to the DMSO control condition (CTRL) from OCSCs shown in Supplementary Figure 3. **E)** Heat map of the p-values of quantifications shown in A. Blue: $p>0.05$. Red: : $p<0.05$. **F)** Histogram of the *in vitro* cell viability of ONS-76 cells after 5 days of treatment (n=3).

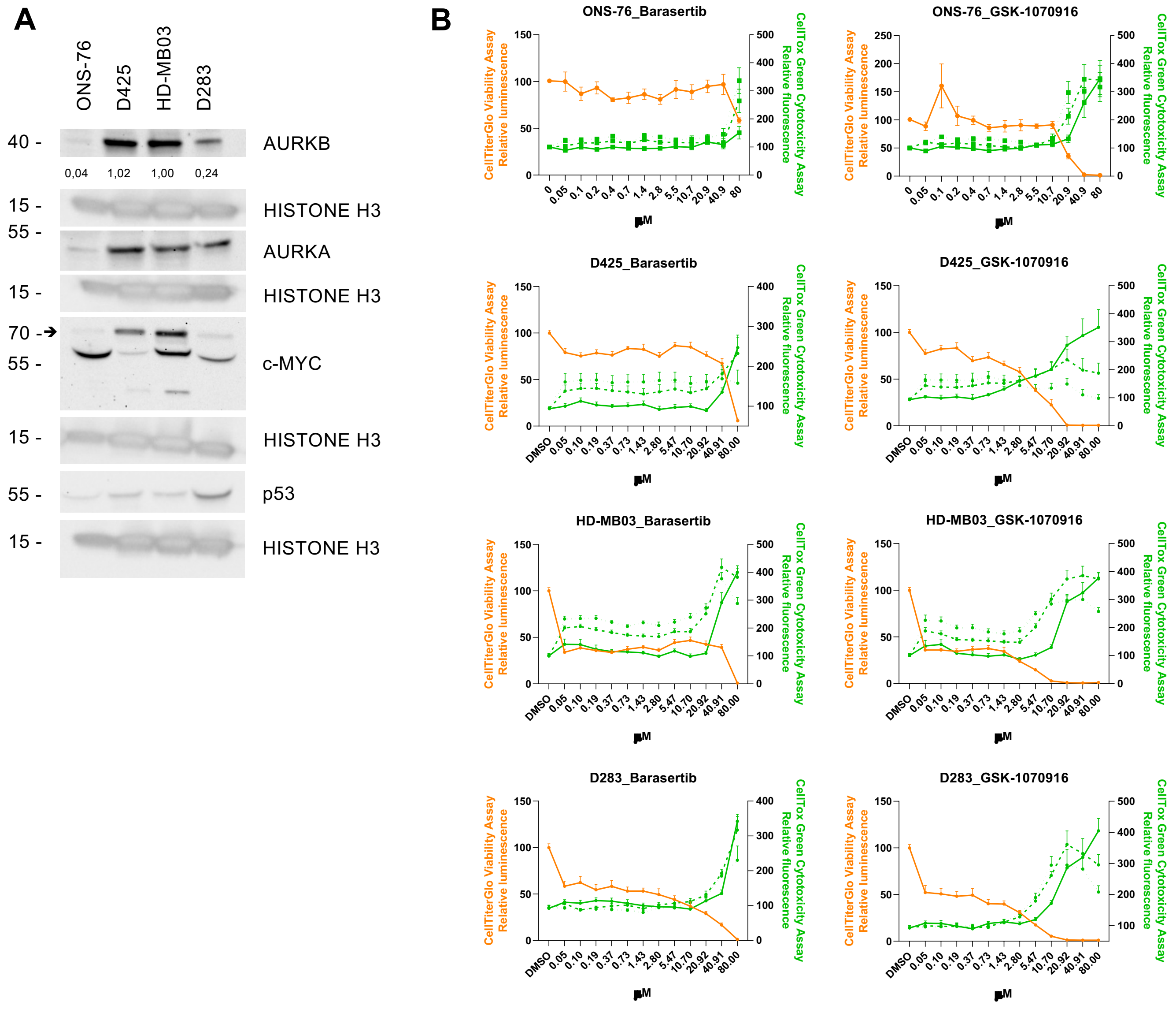


**Supplementary Figure 5: AURKAi and AURKBi drug response profiling**

**A)** Western Blot of basal protein expression levels in ONS-76, D425, HD-MB03 and D283 MB cell lines using antibodies against AURKA, AURKB, c-MYC, p53 and histone H3 as control. **B)** Cell viability (CellTiterGlo 3D (orange)) and cell toxicity (CellTox™ Green Dye (green)) analysis on MB cells exposed to increasing concentrations of Barasertib or GSK-1070916 in 3D suspension cultures. The lumiescence and fluorescence levels was measured after 24h (a, full), 48h (b, dashes) or 72h (c, dotted lines). n=3 technical replicas, mean + SEM are shown.

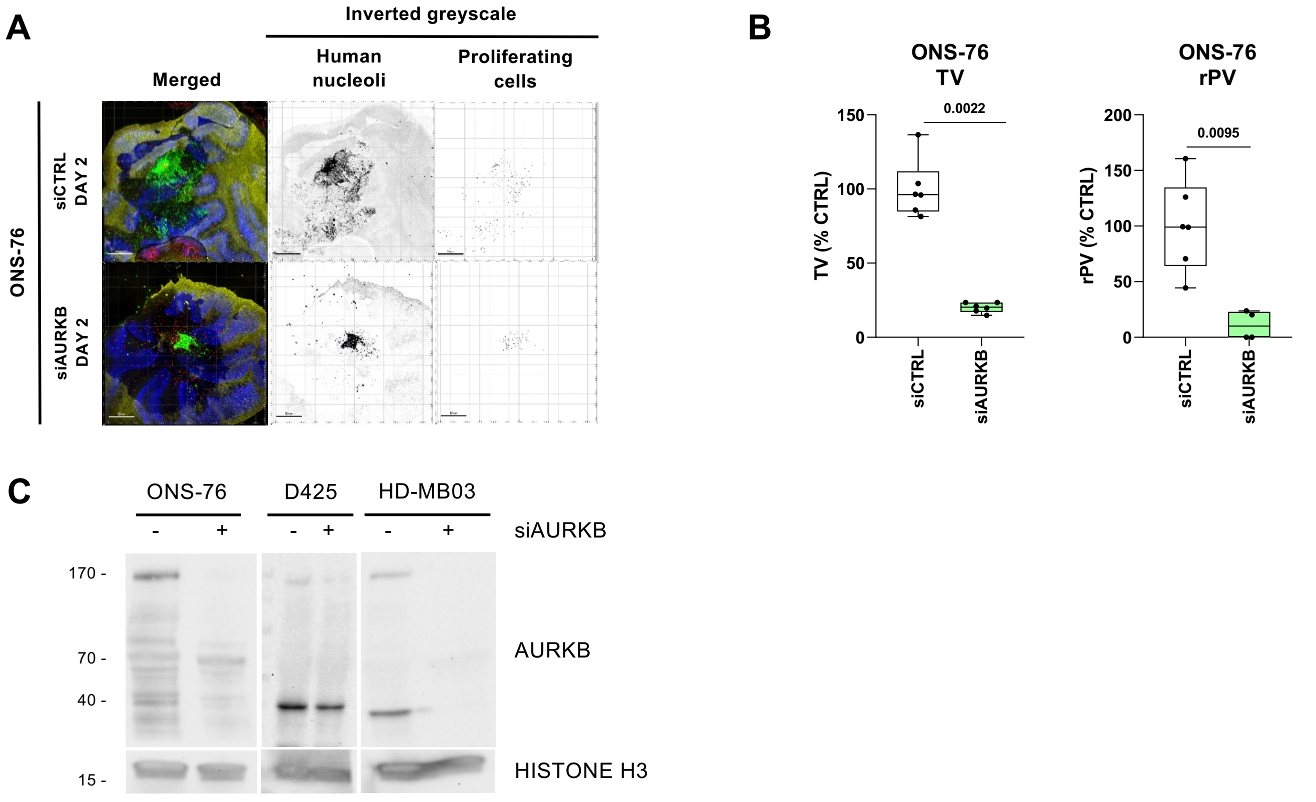


**Supplementary Figure 6: siRNA-mediated depletion of AURKB represses tumor cell growth in the tissue**

**A)** Confocal microscopy analysis of OCSCs with siCTL or siAURKB-transfected ONS-76 cells (green). Red: Click-iT-EdU, blue: anti-Calbindin (Purkinje cells), yellow: anti-GFAP. **B)** siAURKB ONS-76 tumor cell volume (TV) and relative proliferation volume (rPV) relative to siRNA scramble control cells (siCTRL). **C)** Western Blot illustrating the cell-specific efficacies of AURKB depletion by siRNA after 72 h of transfection.

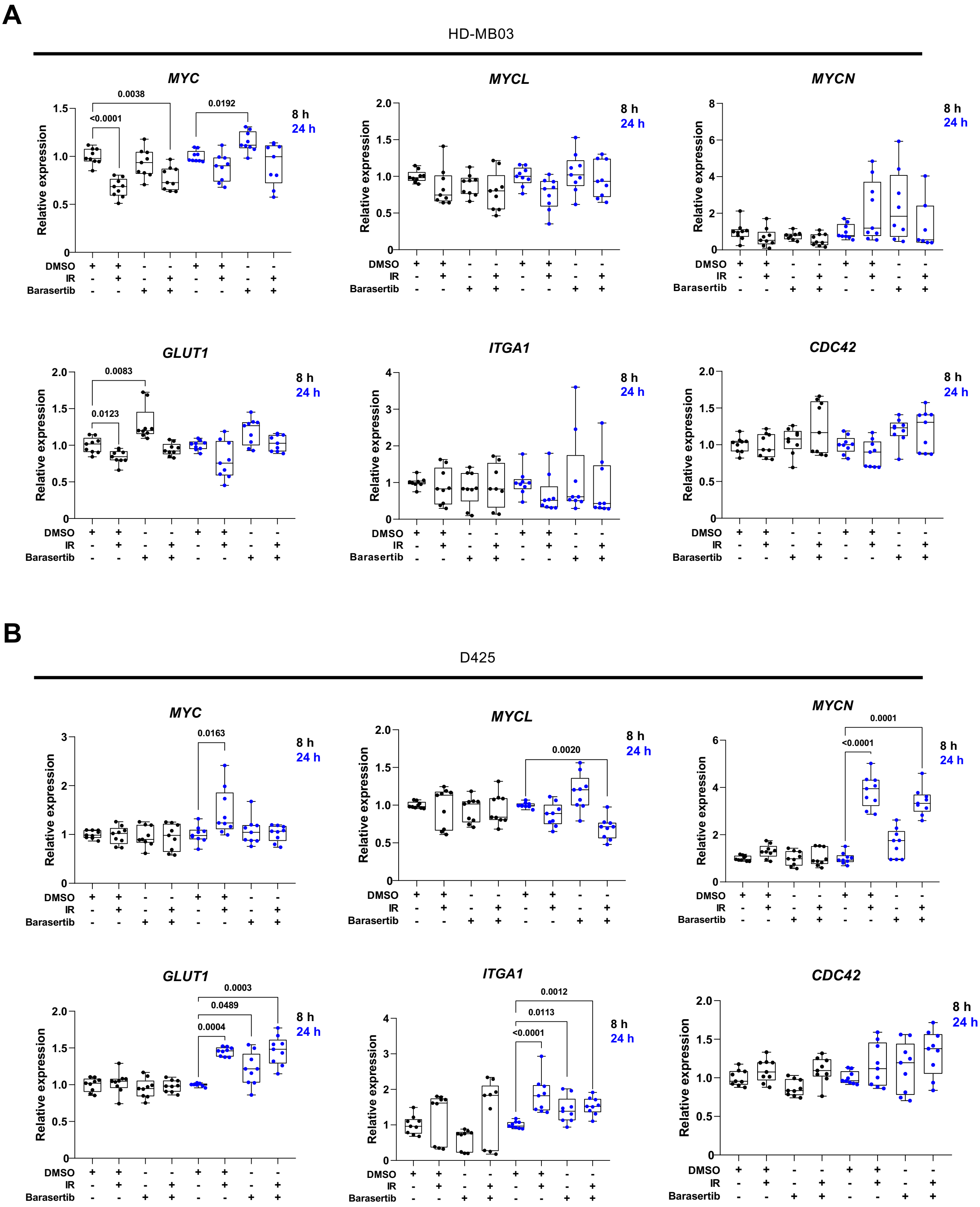


**Supplementary Figure 7:AURKB inhibitor Barasertib does not interfere MYC family and MYC target gene expression**

RT-qPCR quantifications of *c-MYC, MYCL, MYCN, GLUT1, ITGA1* and *CDC42* mRNA levels in HD-MB03 **(A)** and D425 **(B)** n=3 biological replicas. Cells were X-ray irradiated (IR; 1,8 Gy) and/or treated with Barasertib (50 nM) for 8 h and 24 h.

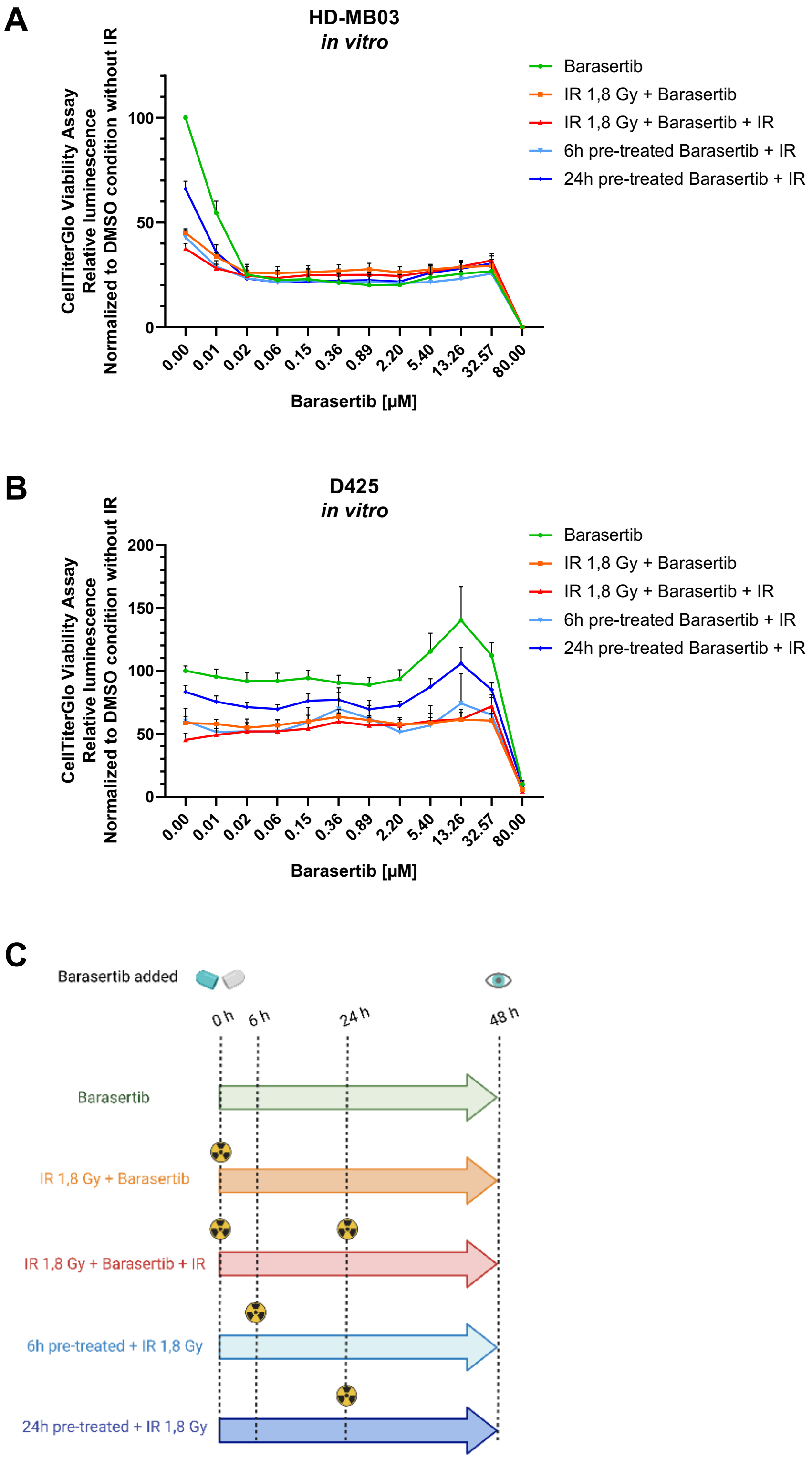


**Supplementary Figure 8: No additive effect on cell viability *in vitro* of Barasertib treatment in combination with radiation. A)** HD-MB03 and **B)** D425 cell viability in response to irradiation (1,8 Gy) and Barasertib treatment alone or in combination after 48 h following the procedures illustrated in **C.** n=3 technical replicas, mean + SEM are shown. Levels are normalized to DMSO condition without irradiation.

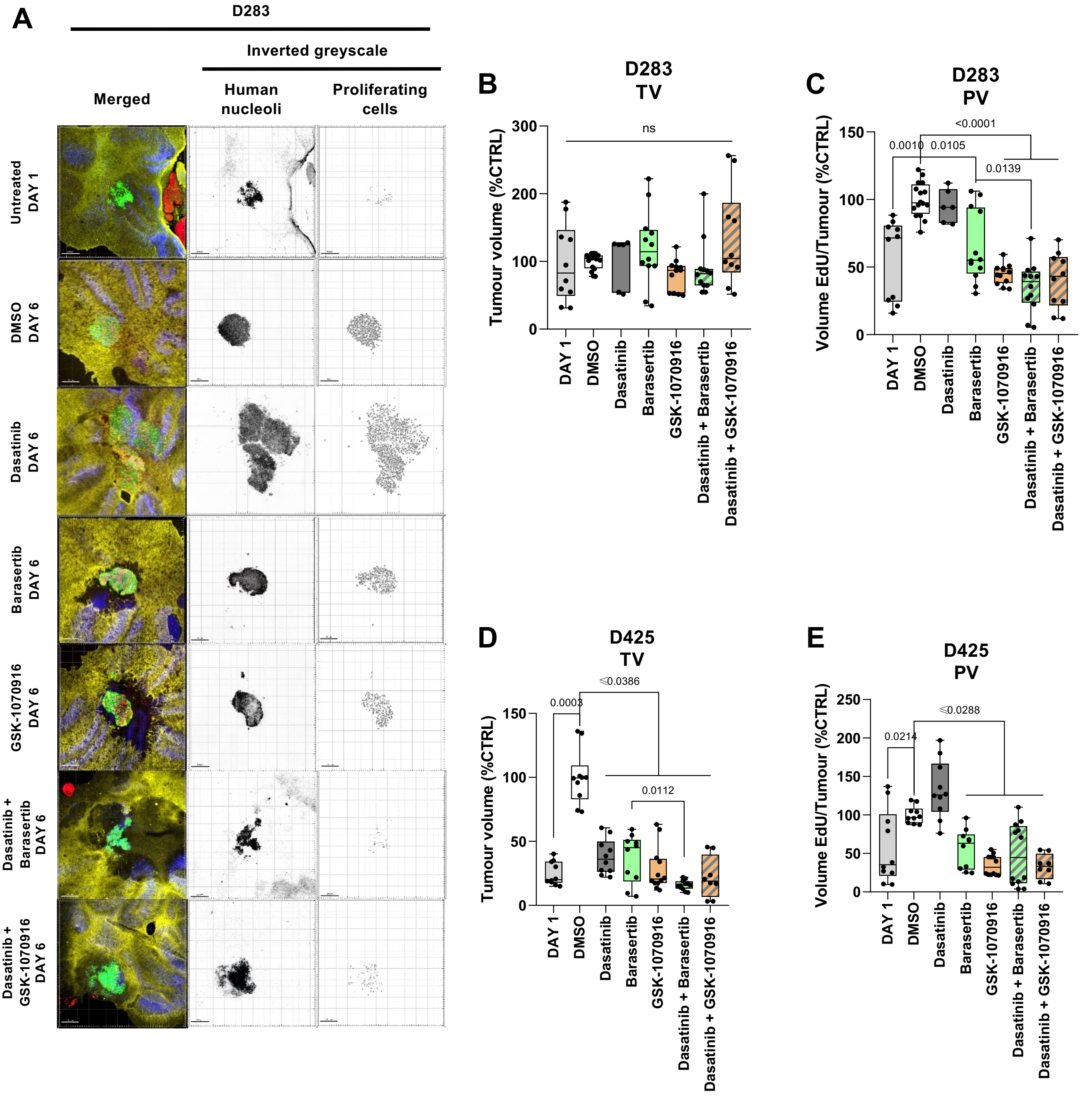


**Supplementary Figure 9: Drug combination *ex vivo* in OCSCs using D283 cell line**

**A)** Confocal analysis of OCSCs 5-days after treatments (100 nM drug concentrations). Green: anti-human nuclei, red: Click-iT EDU, blue: anti-calbindin, yellow; anti-GFAP. **B-E)** Tumour cell volume (TV) and relative proliferation volume (rPV) relative to the DMSO control condition (CTRL) of D283 and D425 cells (representative pictures are shown in Figure 5) embedded in OCSCs.

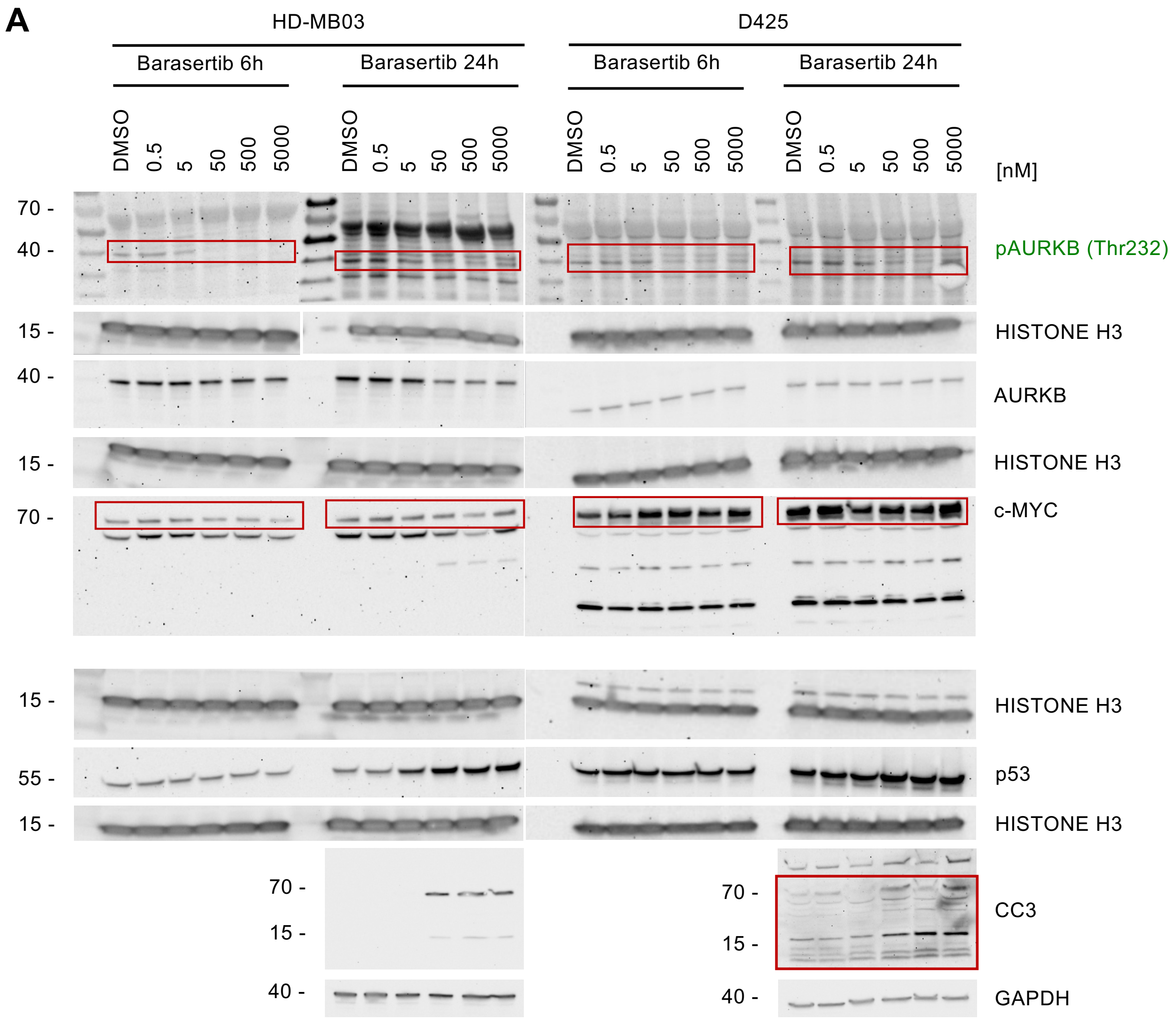


**Supplementary Figure: Western Blot raw data**
